# A minimalistic resource allocation model to explain ubiquitous increase in protein expression with growth rate

**DOI:** 10.1101/015180

**Authors:** Uri Barenholz, Leeat Keren, Eran Segal, Ron Milo

## Abstract

Most proteins show changes in level across growth conditions. Many of these changes seem to be coordinated with the specific growth rate rather than the growth environment or the protein function. Although cellular growth rates, gene expression levels and gene regulation have been at the center of biological research for decades, there are only a few models giving a base line prediction of the dependence of the proteome fraction occupied by a gene with the specific growth rate.

We present a simple model that predicts a widely coordinated increase in the fraction of many proteins out of the proteome, proportionally with the growth rate. The model reveals how passive redistribution of resources, due to active regulation of only a few proteins, can have proteome wide effects that are quantitatively predictable. Our model provides a potential explanation for why and how such a coordinated response of a large fraction of the proteome to the specific growth rate arises under different environmental conditions. The simplicity of our model can also be useful by serving as a baseline null hypothesis in the search for active regulation. We exemplify the usage of the model by analyzing the relationship between growth rate and proteome composition for the model microorganism *E.coli* as reflected in two recent proteomics data sets spanning various growth conditions. We find that the fraction out of the proteome of a large number of proteins, and from different cellular processes, increases proportionally with the growth rate. Notably, ribosomal proteins, which have been previously reported to increase in fraction with growth rate, are only a small part of this group of proteins. We suggest that, although the fractions of many proteins change with the growth rate, such changes could be part of a global effect, not requiring specific cellular control mechanisms.

## 1 Introduction

Many aspects of the physiology of microorganisms change as a function of the growth environment they face. A fundamental system biology challenge is to predict and understand such changes, specifically, modifications in gene expression as a function of the growth environment.

Early on it was found that the expression of some genes is coordinated with growth rate, rather than with the specific environment. Classic experiments in bacteria have shown that ribosome concentration increases in proportion to the specific growth rate [34]. The observed increase in concentration has been interpreted as an increased need for ribosomes at faster growth rates [25, 9, 43, 23]. The search for underlying mechanisms in *E.coli* yielded several candidates such as the pools of ppGpp and iNTP [24, 2], and the tRNA pools through the stringent response [7, 3]. For a more thorough review see [26].

In the last two decades, with the ability to measure genome-wide expression levels, it was found that changes in gene expression as a function of growth rate are not limited to ribosomal genes. In *E.coli,* the expression of catabolic and anabolic genes is coordinated with growth rate, and suggested to be mediated by cAMP [32, 42, 30]. In *S.cerevisiae,* it was shown that most of the genome changes its expression levels in response to environmental conditions in a manner strongly correlated with growth rate [16, 4, 6, 11]. Studies examining the interplay between global and specific modes of regulation, suggested that global factors play a major role in determining the expression levels of genes [10, 19, 18, 37, 1, 16, 11, 40, 12]. In *E.coli,* this was mechanistically attributed to changes in the pool of RNA polymerase core and sigma factors [17]. In *S.cerevisiae,* it was suggested that differences in histone modifications around the replication origins [31] or translation rates [4] across conditions may underlie the same phenomenon. Important advancements in *E.coli* were achieved by analyzing measurements of fluorescent reporters through a simplified model of gene expression built upon the empirical scaling with growth rate of different cell parameters (such as gene dosage, transcription rate and cell size)[19]. Taken together, these studies suggest that the expression of all genes changes with growth rate, with different factors and architectures of regulatory networks yielding differences in the direction and magnitude of these changes [19, 18].

Despite these advancements, many gaps remain in our understanding of the connection between gene expression and growth rate, primarily regarding the underlying mechanisms. Are there unique factors controlling specific groups of genes, as is suggested by [42, 30, 12, 2] and others, or is there a more global phenomenon shared across most genes in the genome? What fraction of the variability observed in gene expression patterns across different growth conditions results from active adaptation to the specific condition? To what extent are large clusters of genes regulated by ’’master regulator” factors such as cAMP, and how much by global, gene and condition-independent, response? Genome-wide proteomic data sets, which take a census of the proteome composition at different growth rates, offer potential insights into these questions and can serve as a basis to explore and compare different models of regulation [40, 35, 12, 30].

In this work we present a parsimonious model that quantitatively predicts the relationship between protein abundance and specific growth rate in the absence of gene-specific changes in regulation. Our model provides a baseline for the behavior of endogenous genes in conditions between which they are not differentially regulated, without the need for condition-specific parameters. The model predicts an increase in protein expression with specific growth rate as an emerging property that is the result of passive redistribution of resources, without need for specific regulation mechanisms. On top of this baseline model, different regulatory aspects can be added. We tested the model against two recently published proteomic data sets of *E.coli* spanning different growth conditions[30, 35]. We find a coordinated, positive correlation between the specific growth rate and the fraction of many proteins, from diverse functional groups, out of the proteome. Although this response accounts for a relatively small part of the total variability of the proteome it is highly relevant for understanding proteome wide studies, as it describes the behavior of over 50% of the proteome genes. The well-studied ribosomal proteins are found to be a small subset of this group of proteins that increase their fraction with the specific growth rate. Our analysis suggests that, even if changes in the proteome composition are complex, for a large number of proteins and under many conditions such changes take the form of a linear, coordinated, increase with growth rate. An increase that can result from cellular resources being freed by down-regulated proteins. The well studied scaling of ribosome concentration with growth rate can be considered one manifestation of this more general phenomena.

## 2 Results

### 2.1 Simple considerations predict passively driven increase in the fraction of proteins as a function of the specific growth rate

What is the simplest way to model the differences in the proteome composition of two populations of cells, one growing in a permissive environment, and the other facing a more challenging growth condition? In an attempt to parsimoniously analyze such differences, we have constructed a minimalistic model that predicts the behavior of non-differentially regulated genes across different growth conditions. Before presenting the model mathematically, we give a brief intuitive depiction.

The model assumes that under a favorable growth condition, the cell actively down-regulates some proteins that are only needed in harsher conditions, as illustrated in Figure 1. The down regulation of the lac operon in the presence of glucose is a prominent example for this phenomenon. As a result of only that specific change, the fraction of all other proteins out of the proteome is increased compared to the harsher (e.g. growth on lactose) condition. In our baseline model, all the other proteins increase their levels and are expected to show the same relative ratios between each other in all conditions. Specifically, the levels of the proteins forming the bio-synthetic machinery increase, increasing the ratio of the bio-synthetic machinery to the proteome. The growth rate is dependent on the amount of protein bio-synthesis a cell performs. The increase in ratio of bio-synthetic machinery to proteome thus results in an expected increase in the growth rate, as depicted in Figure 1. In our example of the lac operon, in the presence of glucose, the down regulation of lactose metabolism genes leads to faster growth as more bio-synthetic genes are expressed instead.

**Figure 1:**
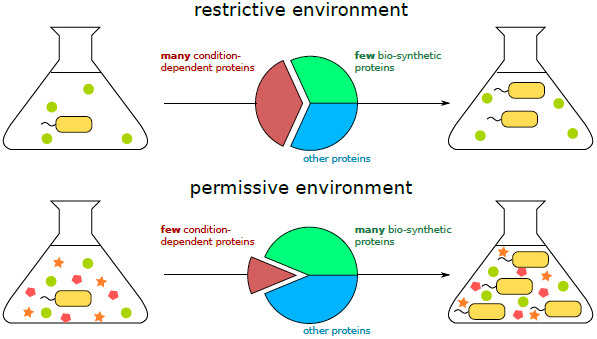
A minimalistic model predicts that low expression of condition-dependent genes under permissive growth environment, compared with a restrictive environment, implies larger fraction of all other proteins out of the proteome. With this, the ratio of bio-synthesis genes to the rest of the proteome is higher in permissive environments, resulting in faster growth.

#### 2.1.1 The expression level of a protein can be decomposed into gene specific control and global expression machinery availability

The composition of the proteome can in principle be determined by a large number of parameters. For example, given that an organism expresses 1000 genes across 10 different growth conditions, one could imagine that controlling the expression pattern of all genes across all conditions will require 10,000 parameters (setting the level of every gene in every condition). Our model proposes an underlying architecture that drastically reduces this amount of parameters, implying that cells control most of the composition of their proteome through fewer degrees of freedom than might be naively expected.

The model separately considers the resulting fraction of every protein out of the proteome as the product of two control mechanisms: (A) Protein/gene specific controls which only affect the individual protein under a given condition. These include the gene associated promoter affinity, 5’-UTRs, ribosomal binding site sequence, as well as the presence of specific transcription/translation factors that react with the relevant gene. We note that while a given transcription factor may affect many genes, the presence or absence of its relevant binding sequence is gene specific, making this control mechanism gene specific in the context of our analysis. While some of these controls (such as the ribosomal binding sites) are static, and therefore condition independent, others are dynamic and will differ across different environmental conditions (such as transcription factors state, for genes that are affected by them). (B) Global expression control based on the availability of bio-synthetic resources, including RNA polymerase, co-factors, ribosomes, amino-acids etc. All of these factors can potentially differ across different environmental conditions and no gene can avoid the consequences of changes in them.

In the model, every gene is given an ‘affinity-for-expression’ (or ‘intrinsic-affinity’) score that encapsulates its tendency to attract the bio-synthetic machinery, as was first suggested in [21]. This gene-specific value can in principle change across conditions but a key feature is that the gene intrinsic affinity tends to have the same value across many conditions. Often two values are enough across all conditions, an “off” and “on” value. We denote the affinity of gene *i* under growth condition *c* by *w_i_*(*c*). To determine the resulting fraction of every protein, our model assumes that the bio-synthetic resources are distributed among the genes according to those affinities, as is stated in Equation 1. Intuitively, one can think of a competition between the genes and transcripts over the bio-synthetic resources, where each gene/transcript attracts resources according to its intrinsic affinity.

The notion of intrinsic affinities represents the expression pattern under a given condition by the intrinsic affinities of all genes under that condition. Assuming each gene has only a finite set of affinities, possibly only one or two (for example, on and off states of the lac operon), describing the expression pattern is therefore reduced to selecting which, out of the total gene-specific small set of possible affinities, each gene gets under the relevant condition. Given that the selection of expression level for a given gene is driven by some specific environmental cues (translated to, for example, activation of specific transcription factors), the description can be further reduced to determining what cues are present at each condition.

The fraction of a specific protein out of the proteome is equal to the specific affinity of the corresponding gene under the condition, divided by the sum of the affinities of all genes under that same condition. To illustrate: if two genes have the same affinity under some condition, their corresponding proteins will occupy identical fractions out of the proteome. If gene *A* has twice the affinity of gene *B* under a given condition, then the fraction protein *A* occupies will be twice as large as the fraction occupied by protein *B* under that condition, etc.

This relationship can be simply formulated as follows:

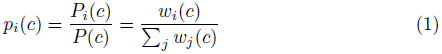
where *p_i_*(*c*) denotes the fraction out of the proteome of protein *i* under condition *c, p_i_*(*c*) denotes the mass of protein *i* under condition *c* per cell, *P*(*c*) denotes the total mass of proteins per cell under condition *c*, and the sum, Σ*_j_ w_j_*(*c*), is taken over the intrinsic affinities of all the genes the cell has.

This equation emphasizes that the observed fraction of a protein is determined by the two factors mentioned above: the specific affinity of the protein/gene, that is present in the numerator, and also, though less intuitive, the affinity of all other genes under the growth condition (affecting the availability of bio-synthetic resources), as reflected by the denominator.

For simplicity, the model refers to the fraction of each specific protein in the proteome and not to the protein concentration. The corresponding concentration in the biomass can be calculated using the concentration of total protein in the biomass. In *E.coli,* this concentration, is known to slightly decrease in a linear manner with the specific growth rate [5, 40, 38] (for further discussion see 4.1.2).

#### 2.1.2 A change in growth condition triggers changes in expression of specific proteins that indirectly affect the whole proteome

Different environmental conditions require the expression of different genes. For example, the expression of amino-acids synthesizing enzymes is required only in culture media lacking amino-acids [20, 39]. Therefore, the cell can infer the presence or absence of amino-acids in the growth media and, regulate the affinities of the synthesizing genes accordingly. If we consider a gene *i*, whose specific affinity is not dependent on the presence of amino-acids, we suggest that its fraction will still change between the two conditions as the affinities of other condition specific genes change, thereby redirecting the bio-synthetic capacity. In mathematical terms this will change the denominator in equation 1 and thus affect the distribution of resources between all of the expressed genes.

Generalizing this notion, we can divide the proteins into those whose intrinsic affinity remains constant across all of the considered conditions, and those whose intrinsic affinity changes between at least some of the conditions (Figure 1). An interesting consequence is that proteins whose intrinsic affinities remain constant also maintain their relative ratios across these conditions with respect to each other, as observed experimentally in *S.cerevisae* in [16].

#### 2.1.3 Growth rate is the combined outcome of proteome composition and environmental conditions

While it is sometimes implied that different cellular components are regulated by the growth rate, our model considers the growth rate as an outcome of the environmental conditions that affect the proteome composition. Specifically, we assume that the doubling time is proportional to the ratio of the total amount of proteins per cell and the proteins involved in bio-synthesis in that cell. The larger the ratio of total proteins to bio-synthesis proteins is, the longer these bio-synthesis proteins will need to duplicate the proteome, resulting in a longer doubling time.

To illustrate this assumption concretely, one could think about the synthesis of polypeptides. If a cell has *R* actively translating ribosomes, each of which synthesizing polypeptides at a rate of *η* ≈ 20 amino acids per second, the biosynthetic capacity of the cell will be limited to ≈ *ηR* amino acids per second. If the total amount of protein in that same cell is *P* (measured in amino acids count), it follows that the time it will take the actively translating ribosomes to synthesize the proteins for an identical daughter cell is 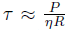 (up to a ln(2) factor resulting from the fact that the ribosomes also synthesize more ribosomes during the replication process and that these new ribosomes will increase the total rate of polypeptides synthesis).

The theoretical lower limit of the doubling time, *T_B_*, will be achieved when all of the proteome of the cell is the bio-synthetic machinery. If the bio-synthetic machinery is only half of the proteome, the doubling time will be *2T_B_* etc.

To integrate the notion of total protein to bio-synthetic protein ratio into our model, we make the following simplifying assumption: There is a group of biosynthetic genes (e.g. genes of the transcriptional and translational machineries) the affinities of which remain constant across different growth conditions, that is, these genes are not actively differentially regulated across different conditions. Furthermore, we assume that the machineries these genes are involved in, operate at relatively constant rates and active to non-active ratios across conditions, as has been shown for ribosomes [5].

Formally, we define the group of bio-synthesis genes, *G_B_*, such that, for every gene that belongs to this group, *k ∈ G_B_*, its affinity, *w_k_*(*c*) is constant regardless of the condition, *c*.

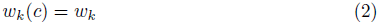

To keep our notations short, we will define the condition independent sum over all of these bio-synthesis genes as the constant:

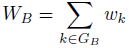

The doubling time under a given condition, *τ*(*c*), will be proportional to the ratio of total protein to bio-synthesis protein under that condition, with the proportionality constant *T_B_*:

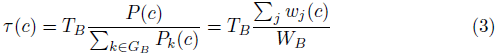

Therefore, the model reproduces an increase in the doubling time for conditions requiring larger amounts of non-bio-synthetic proteins (i.e. higher values in the sum across *w_j_*).

#### 2.1.4 The fraction of a non-differentially regulated protein is expected to increase with the growth rate

Recalling that the connection between the growth rate and the doubling time is: 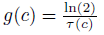, we now combine Equation 1 with Equation 3 to get a prediction for the single protein fractions *p_i_*:

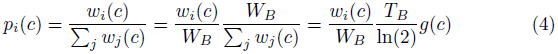

By incorporating all the condition-independent constants (*W_b_, T_b_*, ln(2)) into one term, A, we can simplify to:

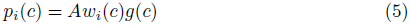

Hence, for every two conditions between which gene *i* maintains its affinity, (*w_j_*(*c_i_*) = (*w_i_*(*c_2_*)), the fraction *p_i_*(*c*) protein *i* occupies in the proteome scales in the same way as the growth rate (*g*(*c*)) between these two conditions.

To summarize, the simplified model we have constructed predicts that, under no specific regulation, the fraction a non-regulated protein occupies out of the proteome should scale with the growth rate. A group of such proteins should therefore maintain their relative ratios across conditions.

#### 2.1.5 Protein degradation differentiates between measured growth rate and biomass synthesis rate

In the following two sections we analyze the effects of expanding our model to account for two biological effects: protein degradation and changes in the rates at which molecular machines operate.

The model we developed predicts that when the growth rate approaches zero, the fraction of every protein with constant affinity also approaches zero. This approach to zero applies specifically to the bio-synthesis genes, that have constant affinities according to our assumptions. However, it is known that the fractions of these proteins, and specifically of ribosomal proteins does not drop to zero when the growth rate approaches zero [14, 29]. We can account for this phenomenon by including protein degradation in our model.

We assume the degradation rate to be constant for all genes and conditions. The *observed* growth rate, *g*, is the amount of proteins produced *minus* the amount of proteins degraded. To illustrate, at zero growth rate, the implication is not that no proteins are produced, but rather that proteins are produced at exactly the same rate as they are degraded.

Integrating this notion into the model means that the bio-synthesis capacity needs to suffice to re-synthesize all the degraded proteins. Hence, where the equations previously referred to the cellular growth rate, *g*, as the indicator of protein synthesis rate, they should in fact refer to the cellular growth rate plus the degradation rate, as that is the actual rate of protein synthesis. If we denote by α the degradation rate, Equation 5 should thus be rewritten as:

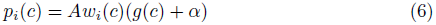

This equation predicts linear dependence of the fraction of unregulated proteins on the growth rate, with an intercept with the horizontal axis occurring at minus the degradation rate (Figure S1). Thus, at zero growth rate, the fraction of non-differentially regulated proteins out of the proteome is positive, equalling *Aw_i_*(*c*)*α*.

#### 2.1.6 Slower biological processes rates at slower growth affect the relation between proteome composition and growth rate

The simplified model assumes that the doubling time is proportional to the ratio of total protein to bio-synthetic protein. This assumption fails if the rate at which each bio-synthetic machine operates changes across conditions. Replacing this assumption by an interdependence of bio-synthesis rate with growth rate (such that, the faster the growth, the faster the synthesis rates, per machine) [5, 40], will affect the resulting predictions as well. This effect is formally analyzed in section 6.1. Slower bio-synthesis rates under slower growth rates imply that, compared with the model prediction, higher fraction of bio-synthesis proteins is needed to achieve a given growth rate. Thus, lower synthesis rates under slower growth rates will be reflected by a lower slope and higher interception point for non-regulated proteins than those predicted by the constant-rate version of the model, as is depicted in Figure S1.

To summarize, our theoretical model predicts that the default behavior of non-differentially regulated proteins between two conditions is to maintain a fraction that is proportional to the growth rate. The faster the growth rate, the higher the fraction. Such proteins should maintain their relative concentrations w.r.t. each other. Degradation and changes in rates of molecular machineries at slow growth result in predicting non-zero fraction for such proteins even when the growth rate is zero, resulting in a more moderate response of the fraction to the growth rate.

### 2.2 Analysis of proteomic data sets

Our theoretical model predicts that the fraction of many proteins proportionally and coordinately scales up with the specific growth rate across different growth conditions. To assess the extent to which this prediction is reflected in actual proteome compositions, we present analysis of two published proteomics data sets of *E.coli,* [30] and [35]. These data sets use mass spectrometry to evaluate the proteomic composition of *E.coli* under 23 different growth rates using an accelerostat [28], and 20 different growth conditions, spanning both different carbon sources and chemostat-controlled growth rates, respectively. The data set from [35] contains more conditions than those analyzed below, see section 4.1.3 for further details.

#### 2.2.1 A large fraction of the proteome is positively correlated with growth rate

Our model predicts that a large portion of the proteins should increase in fraction with the growth rate, as that is the expected change for proteins that are not specifically regulated between conditions. To test this prediction, we calculated the Pearson correlation of every protein with the growth rate (Figure 2, upper panels). We find that about a third of the proteins (473 out of 1442 measured in the data set from [35], and 305 out of 1142 in the data set from [30]) have a strong positive (> 0.5, see also 6.2) correlation with the growth rate. These values are much higher than those obtained for randomized data sets (12 and 5 strongly positively correlated proteins for the two data sets, respectively, as is further discussed in section 2.2.4 and is seen in Figure 2 lower panels). Strong negative correlation with growth rate is much less common in the data set from [35]. It is common in the data set from [30], where we speculate that it results from the specific way by which growth rate was controlled, namely by implicitly controlling nutrient concentration via an accelero-stat. The control of growth rate by gradual changes to nutrient concentration may naturally lead to gradual changes in protein levels, both increasing and decreasing, an effect that is expected under any regulatory scheme and is thus irrelevant for our analysis. Notably, in both data sets, the proteins that have a high correlation with the growth rate are involved in many and varied cellular functions and span different functional groups (See tables S1 and S2).

**Figure 2:**
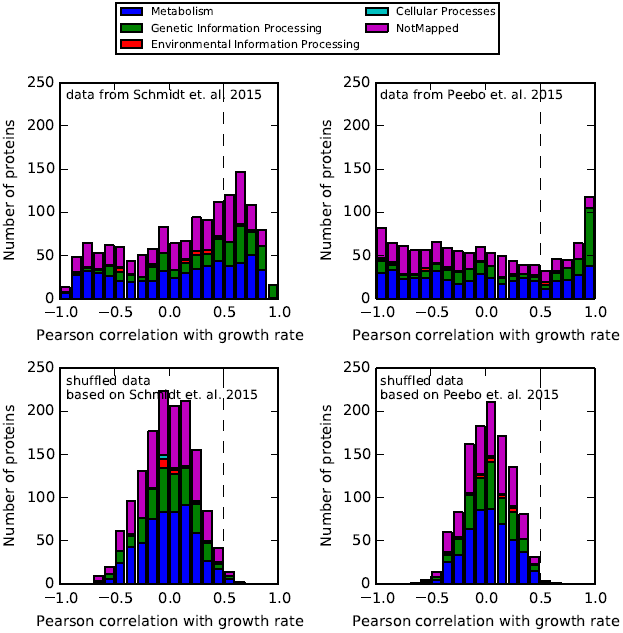
A strong positive Pearson correlation between the fraction out of the proteome and the growth rate is observed for a large number of proteins in two data sets (upper panels). Functional protein groups are denoted by different colors. Thresholds defining high correlation are marked in dashed lines and further discussed in 6.2. Shuffling the amounts of every protein across conditions reveals the bias towards positive correlation with growth rate is nontrivial (lower panels).

Previous studies already found that ribosomal proteins are strongly positively correlated with growth rate [29, 14, 17]. Our analysis agrees with these findings as we find the fraction of the vast majority of the ribosomal proteins to be strongly positively correlated with growth rate (47 out of 53 in the data set from [35] and 52 out of 53 in the data set from [30]). However, we also find that the group of proteins strongly positively correlated with growth rate reach far beyond the previously discussed group of ribosomal proteins (tables S1 and S2). Importantly, the proteins that we find to be strongly positively correlated with growth rate are not generally expected to be co-regulated, and their behavior does not seem to be the result of any known transcription factor or regulation cluster response [33].

#### 2.2.2 Proteins positively correlated with growth rate share a similar response

Our model predicts that non-differentially regulated proteins should preserve their relative ratios across conditions. We refer to such proteins as being *coordinated* or *coordinately regulated.* We have shown above that many proteins are positively correlated with growth rate. However, we note that having similar correlation with growth rate for different proteins does not imply that such proteins are coordinated, i.e. that they share the same scaling with growth rate. Theoretically, proteins with identical correlation with growth rate may have very different slopes or fold changes with increasing growth rate.

In order to examine how similar the behavior with growth rate is for the group of strongly positively correlated proteins, we normalized each of them to its mean abundance (see 4.1.1) and calculated the slope of a linear regression line for the normalized fraction vs. the growth rate (Figure 3). The slopes of 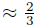 of the proteins lie in the range (0.5, 2) with the highest slopes being ≈ 5. A slope of 0.5 means that the fraction of the protein changes by ±12% around its average fraction in the range of growth rates measured, whereas a slope of 2 indicates a change of ±50%. Hence, the relative amounts of proteins with slopes in the range of (0.5, 2) change by at most just over 2-fold over the range of growth rates measured.

**Figure 3:**
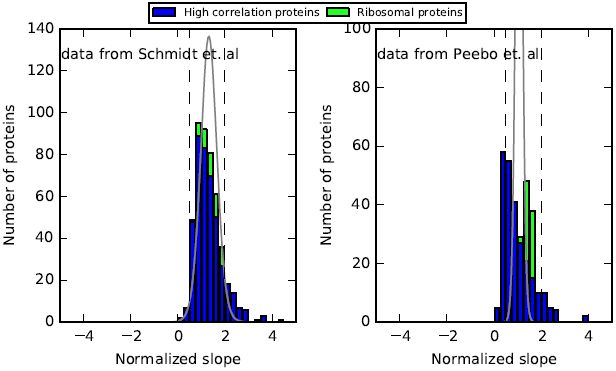
Histogram of the slopes of regression lines for the highly correlated with growth proteins (473 and 305 proteins in the left and right panels respectively). Ribosomal proteins are stacked in green on top of the non ribosomal proteins, marked in blue. Proteins fractions were normalized to account for differences in slopes resulting from differing average fractions (Section 4.1.1). The expected distribution of slopes given the individual deviations of every protein from a linear regression line, assuming all proteins are coordinated, is plotted in gray. Dashed vertical lines at 0.5 and 2 represent the range at which the slopes of 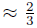 of the proteins lie. Left panel - data from [35], right panel - data from [30]. High correlation proteins share similar normalized slopes, implying they are coordinated, maintaining their relative ratios across conditions (see text for further details). Ribosomal proteins, shown in green, scale with growth rate in a manner similar to the rest of the high correlation proteins (see text and Figure S5).

To understand whether the observed distribution is coordinated, and can result from the noise levels present in the data, we calculated, for every protein, the standard error with respect to the regression line that best fits its fractions. Given these standard errors we generated the expected distribution of slopes that would result by conducting our analysis on proteins that share a single, identical slope, but with the calculated noise in measurement. The expected distribution is shown in gray line in Figure 3 (Further details on the calculation as well as the deviation in maxima between the expected and observed distributions are discussed in the SI). The two data sets show different characteristics of the expected distribution. While the expected distribution corresponding to the data set from [35] coincides with the observed variability in calculated slopes, supporting the notion of a coordinated response, for the data from [30] the expected distribution is much narrower, suggesting a bi-modal distribution. Future studies may uncover the factors underlying the difference between the distributions of the two data sets.

Next we examined how the response of the strongly correlated proteins relates to the well-studied response of ribosomes concentration. To that end, we performed the same analysis of slopes, restricting it to ribosomal proteins alone, as is shown by the stacked green bars in Figure 3. We find that strongly correlated proteins and ribosomal proteins scale in similar ways (slope of 1.37 with *R*^2^ = 0.89 for the sum of ribosomal proteins vs. 1.24 and *R*^2^ = 0.91 for the sum of all strongly correlated proteins, in the data from [35], and slope of 1.49 with *R*^2^ = 0.97 for ribosomal proteins vs. 1.0 and *R*^2^ = 0.97 for all strongly correlated proteins, in the data from [30]. See also Figure S5), implying that the observed response of ribosomal proteins to growth rate is not unique and is coordinated with a much larger fraction of the proteome, thus encompassing many more cellular components.

Our results, showing that a large number of proteins maintain their relative concentrations across different growth conditions thus extend the scope of similar results obtained for *S.cerevisiae* in [16] and for expression levels in *E.coli* under stress conditions [15]. In contrast to other approaches, our model suggests a mechanism for this coordinated expression changes that is not based on shared transcription factors but rather is a result of passive redistribution of resources.

#### 2.2.3 Changes in the proteome across environmental conditions are dominated by proteins that are positively correlated with growth rate

To assess the significance of the positive correlation of proteins with growth rate, out of the total change in proteome composition across conditions, we summed the fractions of all of the proteins that are strongly correlated with growth rate across the conditions measured and plotted their total fraction against the growth rate in Figure 4. Both data sets show that the fraction of these proteins change ≈ 2 fold across a ≈ 5 fold change in the growth rate under the different growth conditions. This change is smaller than the 1: 1 change predicted by our basic model and the deviations may result from the effects of degradation and varying bio-synthesis rates, as is discussed in sections 2.1.5 and 2.1.6. Most of the variability of the total fraction of these proteins can be explained by the growth rate (*R*^2^ of 0.91 in the data set from [35] and 0.97 in the data set from [30]). Importantly, the strongly correlated proteins form a large fraction of the proteome, exceeding 50% of the proteome by weight, at the higher growth rates. This is a much higher fraction than the one obtained for randomized data sets (< 4%, as is further discussed in section 2.2.4) Thus, when considering the changes in proteome composition across conditions, we find that, at higher growth rates, more than 50% of the proteome composition is affected by the coordinated response of the same group of proteins with growth rate.

**Figure 4:**
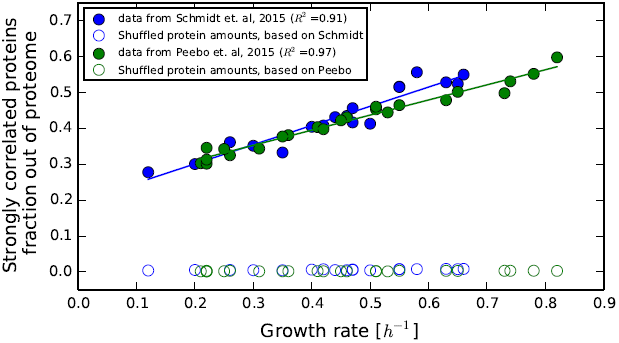
Fraction of the proteome occupied by proteins that are strongly positively correlated with growth rate. The accumulated sum of the proteins that are strongly positively correlated with growth rate (defined as having a correlation above 0.5), as a fraction out of the proteome, with linear regression lines is shown. These proteins form a large fraction (≥ 50%) out of the proteome at higher growth rates. The accumulated fraction of the strongly correlated proteins doubles as the growth rate changes by about 5-fold. Assuming constant degradation rates, the trend lines correspond to protein half life times of ≈ 1.7 hours. Randomized data sets result in much fewer strongly positively correlated with growth rate proteins, implying a much smaller accumulated fraction (hollow circles) as is further discussed in section 2.2.4

Despite the magnitude of this phenomena, the fraction of the total variability in the proteome that is accounted for by this linear response is only ≈ 8% in the data set from [35] and even lower in [30] (Figure S2). While this fraction is low, it is still much higher than the equivalent 2% obtained for a randomized data set based on the data from [35], as is described in section 2.2.4. This relatively low explained variability fraction is primarily the result of two factors: the linear response applies only to < 0.4 of the proteins, leaving the rest of the proteins with no prediction, and experimental noise in whole proteome measurement techniques, estimated at ≈ 20%. Further discussion of the fraction of variability explained can be found in 6.2.

#### 2.2.4 The statistical features we find do not naturally rise in randomized data sets

We performed two tests to verify that the trends we find, namely, the large fraction of proteins with a strong correlation with growth rate, the coordination among these proteins, their large accumulated fraction out of the proteome, and the fraction of variability explained by a single linear regression approximation of their fractions, are all non-trivial characteristics of the data set that do not naturally rise in randomly generated data but that do arise if our model is correct. To this extent we repeated our analysis on two simulated data sets:

- A data set at which the amount of every protein was shuffled across the different conditions.
- A synthetic, simulated, data set, based on the conditions and growth rates of the data set from [35], assuming half the proteins being perfectly coordinated and linearly dependent on growth rate, with parameters similar to those found in our analysis, and the other half having no correlation with growth rate, and with a simulated normally distributed measurement noise of 25%.

We find that in the shuffled sets the number of proteins being significantly positively correlated with growth rate is much smaller than found in the real data sets (12 vs. 473 in the data set from [35] and 5 vs. 305 in the data set from [30]) as is shown in Figure 2, lower panels. As a consequence, these proteins now occupy a much smaller fraction out of the proteome mass-wise (< 4% on average across conditions vs. ≈ 40% in the real data sets) as is shown in Figure 4. Finally, the fraction of variability in the proteome that can be explained by a single linear regression to these proteins is smaller for the shuffled data sets than that obtained for the real data set (2% vs. 8% for a threshold of *R* ≥ 0.5 for the data from [35] and 1% vs. 3.5% for the data set from [30]), as is seen in Figure S6.

We find that the simulated (second) set does display similar characteristics to those we find in the real data, confirming that if, indeed, our model is valid, experimental measurements would overlap with those that we obtained as is seen in Figure S7.

## 3 Discussion

We presented a parsimonious model connecting the fraction of proteins out of the proteome and the growth rate as an outcome of the limited bio-synthesis resources of cells. The notion of intrinsic affinity for expression, first presented in [21], and rarely used ever since, was re-introduced as a key determinant for the differences in expression of different proteins under a given growth condition. The integration of the notion of intrinsic affinity for expression with the limited bio-synthesis capacity of cells was shown to result in a simple mechanism predicting increased fraction of many proteins with the growth rate, without assuming regulation by specific transcription factors for these proteins.

The framework we present emphasizes the importance of accounting for global factors, that are reflected in the growth rate, when analyzing gene expression and proteomics data, as was noted before [21, 4, 19, 18, 37, 1, 16, 11, 40, 12, 30, 41]. Specifically, we suggest that the default response of a protein (that is, the change in the observed expression of a protein, given that no specific regulation was applied to it) is to linearly increase with growth rate. We point out that, as non-differentially regulated proteins maintain their relative abundances, one can deduce the parameters of the linear increase with growth rate of any non-differentially regulated protein by observing the scaling of other such proteins and fixing the ratio between the protein of interest and the reference proteins.

We analyze two recent whole proteome data sets to explore the scope and validity of our model. We characterize a coordinated response in *E.coli* between many proteins and the specific growth rate. This response spans proteins from various functional groups and is not related to the specific medium of growth. A similar phenomena is observed for *S.cerevisiae* as was reported in [16] and may thus be conserved across various organisms and domains of life. Our analysis suggests that, while changes in the proteome composition may seem complex, for a large number of proteins and under many conditions, they can be attributed to a linear, coordinated, increase with growth rate, at the expense of other, down-regulated proteins. The well studied scaling of ribosomes concentration with growth rate can be considered one manifestation of the more general phenomena we describe here. We find that this response is not unique to ribosomal proteins but is, in fact, shared with many other proteins spanning different functional groups. Furthermore, the linear dependence slope and explained variability of fraction levels of proteins explained by linear correlation with growth rate is similar among the ribosomal proteins versus all the proteins with high correlation with the growth rate.

Many studies monitored the ribosome concentration in cells and its interdependence with growth rate [34, 5, 43, 37, 40, 30, 12](many of them indirectly). While in all of these studies a linear dependence of ribosome concentration with growth rate was observed, in some cases different slopes and interception points were found to describe this linear dependence, compared with the observations in our study. A discussion of various reasons that may underlie these differences is given in section 6.4.

Interestingly, our model suggests that a linear correlation between ribosomal proteins and the growth rate might be achieved without special control mechanisms. Nonetheless, many such mechanisms have been shown to exist [26, 36]. We stress that the existence of such mechanisms does not contradict the model. Mechanisms for ribosomal proteins expression control may still be needed to achieve faster response under changing environmental conditions or a tighter regulation to avoid unnecessary production and reduce translational noise. Furthermore, such mechanisms may be crucial for synchronizing the amount of rRNA with ribosomal proteins as the two go through different bio-synthesis pathways. Nevertheless, the fact that many non-ribosomal proteins share the same response as ribosomal proteins do, poses interesting questions regarding the scope of such control mechanisms, their necessity and the trade-offs involved in their deployment.

The findings in this study support and broaden the findings in other recent studies. Specifically, for *S.cerevisiae* a few recent studies found that the concentration of the majority of the proteins is coordinated across conditions and increases with growth rate [16, 10, 3]. In principle, the model we suggest here can be applied to any exponentially growing population of cells and may thus also serve as a potential explanation for the phenomena observed in these studies and others.

Other recently published studies in *E.coli* have suggested different models and in some cases have results and predictions that do not coincide with those presented in this study. Notably, in [19, 18] a decreased protein concentration for unregulated genes is predicted. A few differences can explain this seeming discrepancy. The modeling in [19] relies on data collected under higher growth rates than those available in the data sets analyzed here. The predictions of the model are based on the deduced dependence of various bio-synthesis process rates and physiological properties of the cells on the growth rate, properties that are, in turn, used to calculate the expected protein concentration for unregulated proteins under the different growth rates. The model in [19] refers to protein concentration and not to protein fraction out of the proteome. These quantities may differ due to changes in the ratio of total protein mass to cell dry weight, or cell volume, as a function of the growth rate. Thus, this approach is markedly different than the approach we take, which assumes relatively small changes in bio-synthetic rates as a function of growth rate and focuses on the limited bio-synthesis resources as the main driver of changes in the resulting fraction of proteins out of the proteome. As the model in [19] was only tested against a handful of proteins, further data collection is required to decide which of the two models better describes the global effects of growth rate on proteome composition.

The expected availability of increasing amounts of whole proteome data sets, with higher accuracy levels, will enable further investigation of the details of cellular resource distribution. With our model serving as a baseline, the analysis of such future data sets will shed more light on the relative roles of carefully tuned response mechanisms versus global, passive effects in shaping the proteome composition under different growth environments.

## 4 Materials and Methods

### 4.1 Data analysis tools

All data analysis was performed using custom written software in the Python programming language. The data analysis source code is available through github at: http://github.com/uriba/proteome-analysis Analysis was done using SciPy [27], NumPy [8] and the Pandas data analysis library [22]. Charts where created using the MatPlotLib plotting library [13].

#### 4.1.1 Normalizing protein fractions across conditions

Our analysis aims at identifying proteins that share similar expression patterns across the different growth conditions. For example, consider two proteins, *A* and *B* measured under two conditions, *c*_1_ and *c*_2_. Assume that the measured fractions out of the proteome of these two proteins under the two conditions were 0.001 and 0.002 for *A* under *c*_1_ and *c*_2_ respectively, and 0.01 and 0.02 for *B* under *c*_1_ and *c*_2_ respectively. These two proteins therefore share identical responses across the two conditions, namely, they double their fraction in the proteome in *c*_2_ compared with *c*_1_.

The normalization procedure scales the data so as to reveal this identity in response. Dividing the fraction of each protein out of the proteome by the average fraction of that protein across conditions yields the normalized response. It the example, the average fraction of A across the different conditions is 0.0015 and the average fraction of B is 0.015. Thus, dividing the fraction of every protein by the average fraction across conditions of that same protein yields:

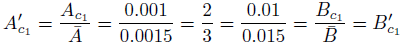
for *c*_1_ and

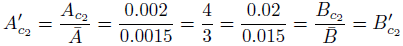
for *c*_2_ showing *A* and *B* share identical responses across *c*_1_ and *c*_2_.

The general normalization procedure thus divides the fraction of protein *i* under condition *c*, *p_i_*(*c*) by the average fraction of protein *i* across all of the conditions in the data set, 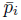, to give the normalized fraction under condition 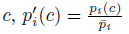

This normalization procedure has been applied prior to calculating the slopes of the regression lines best describing the change in fraction out of the proteome of every protein as a function of the growth rate. Furthermore, when analyzing the variability explained by linear regression on the sum of fractions of all proteins presenting a high correlation with the growth rate, the same normalization procedure was made in order to avoid domination by the high abundance of a few proteins in that group.

#### 4.1.2 Calculation of protein concentration

In this study, we use the mass ratio of a specific protein to the mass of the entire proteome, per cell, as our basic measure for the bio-synthetic resources a specific protein consumes out of the bio-synthetic capacity of the cell. We find this measure to be the best representation of the meaning of a fraction a protein occupies out of the proteome. However, we note that if initiation rates are limiting (e.g. if RNA polymerase rather than ribosomes become limiting), and not elongation rates, then using molecule counts ratios (the number of molecules of a specific protein divided by the total number of protein molecules in a cell) rather than mass ratios may be a better metric. We compared these two metrics and, while they present some differences in the analysis, they do not qualitatively alter the observed results.

There are different, alternative ways to assess the resources consumed by a specific protein out of the resources available in the cell. On top of the measures listed above, one could consider either the total mass or molecule count of a specific protein out of the biomass, rather than the proteome, or out of the dry weight of the cell, both of which vary with the ratio of total protein to biomass or dry weight which was neglected in our analysis. Moreover, one can consider specific protein mass or molecule count per cell, thus reflecting changes in cell size across conditions. Our analysis focuses on the relations between different proteins and resource distribution inside the proteome, and thus avoids such metrics.

#### 4.1.3 Filtering out conditions from the Heinemann data set

The [35] data set contains proteomic data measurements under 22 different environmental conditions. However, our model assumes exponential growth, implying that measurements taken at stationary phase are expected to differ from simple extrapolation of the model to zero growth rate. Therefore, the two measurements of stationary phase proteomics were excluded from our analysis.

Out of the conditions measured in the [35] data set, two conditions included amino acids in the media and presented much faster growth rate than the rest of the conditions (growth in LB media and in glycerol supplemented with AA, with growth rates of 1.9[*h*^−1^] and 1.27[*h*^−1^] respectively, compared with a range of 0.12 — 0.66 for the other conditions). This asymmetry in the distribution of growth rates caused inclusion of these conditions to dominate the analysis due to its effect on the skewness of the distribution of growth rates (γ_1_ = −0.5 for the growth rates excluding LB and AA supplemented glycerol vs. γ_1_ = 2.3 with LB and AA supplemented glycerol) reducing the statistical power of the other conditions. While including the data on growth in these conditions does not qualitatively change the observed results, such analysis is much less statistically robust. We have therefore omitted growth in LB and in AA supplemented glycerol in the main analysis. We present the analysis including these conditions in section 6.3.

## 5 Acknowledgments

We kindly thank Matthias Heinemann for making proteomics data available before publication. We thank Uri Alon, Stephan Klumpp, Arren Bar Even, Avi Flamholz, Elad Noor, Kaspar Valgepea, Karl Peebo and Katja Tummler for fruitful discussions and enlightening comments on this work. This work was funded by the European Research Council (Project SYMPAC 260392); Dana and Yossie Hollander; Helmsley Charitable Foundation; Israel Ministry of Science; The Larson Charitable Foundation. R.M. is the Charles and Louise Gartner professional chair and an EMBO young investigator program member.

## 6 Supplementary figures and data

### 6.1 Effects of degradation and varying synthesis rate on model predictions

The predicted fraction of an unregulated protein as a function of the growth rate is to follow a linear trend crossing at the origin (Figure S1, blue line). Degradation can be interpreted as implying that the observed growth rate is the combination of the bio-synthesis rate minus the degradation rate, implying that the predicted fraction of an unregulated protein is linear increase with growth rate, but with a horizontal intercept at minus the degradation rate as is shown by the green line in Figure S1.

Non constant bio-synthesis rate can be modeled as a Michaelis-Menten kinetic like interdependence with growth rate following the formula:

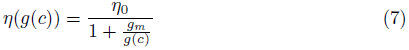
where *g*(*c*) is the growth rate, *η*(*g*(*c*)) is the bio-synthesis rate at growth rate *g*(*c*), *η*_0_ is the maximal bio-synthesis rate and *g_m_* the growth rate at which the bio-synthesis rate is 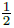 the maximal rate (Figure S1, red line). Under this assumption, the doubling time of the bio-synthesis machinery itself, *T_B_* becomes:

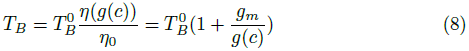
where 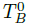 is the minimal theoretical doubling time when all the proteins are bio-synthesis proteins operating at maximal rate. Substituting equation 8 into equation 4 results in a predicted fraction of:

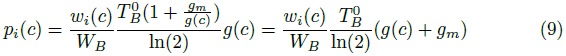

Surprisingly, this equation also describes the fraction as being linearly dependent on the growth rate, with the kinetic parameters implying a non-zero fraction at zero growth rate as is shown by the red dots in Figure S1.

**Figure S1:**
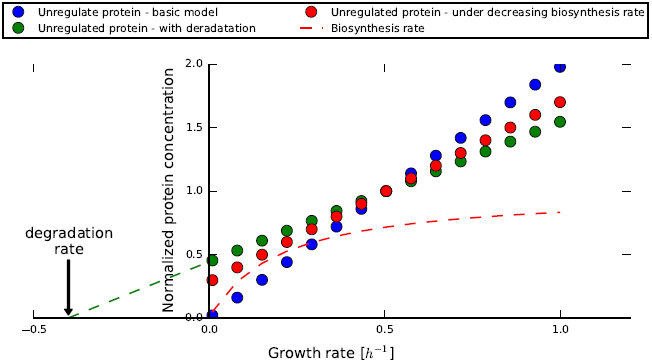
The predictions of the model for the fraction of unregulated proteins as a function of the growth rate. The effects of accounting for protein degradation (green) and Michaelis-Menten like dependence of bio-synthesis rates on growth rate (red) are shown. For non-constant bio-synthesis rate, a growth rate of 0.2 was selected as the growth rate at which the bio-synthesis rate is half of its maximal value.

### 6.2 Threshold selection for defining strong correlation with growth rate

The data we use includes the fractions of proteins under different growth conditions, and the growth rate for every condition. We select a threshold correlation with growth rate to define the group of highly positively correlated with growth rate proteins.

We calculate the explained variability by the growth rate, given a threshold, by taking the difference between the total variability of the group of proteins with a correlation higher than the threshold, and the variability remaining, when assuming these proteins scale with the growth rate according to the calculated linear response. Dividing the explained variability by the total variability of the entire data set quantifies what fraction of the total variability in the proteome is explained by considering a coordinated linear scaling with growth rate for all the proteins with a correlation with growth rate higher than the threshold.

The choice of threshold is thus influenced by two contradicting factors. Choosing a low threshold results in defining many proteins as being highly positively correlated with growth rate. In this case, the correlation with growth rate of these proteins spans a large range. Therefore, applying a linear regression trend to the sum of these proteins only accounts for a small fraction of the variability of them and, as a consequence, only accounts for a small fraction of the total variability of the proteome.

On the other hand, choosing a high correlation threshold results in defining only a small number of proteins as being highly positively correlated with growth rate. A common linear regression line may thus explain a large fraction of the variability for the chosen proteins but, as their number is small, will only account for a small fraction of the total variability of the proteome.

For simplicity, we chose a threshold value of 0.5 for the two data sets analyzed in this study. Figure S2 shows how the choice of threshold affects the fraction of explained variability in the proteome by the linear dependence on growth rate of the proteins that have a correlation with growth rate that is higher than the threshold (blue line). The figure also shows the fraction of proteins that have a correlation with growth rate that is higher than the threshold out of the proteome (red line), and the fraction of explained variability by linear regression for these proteins (green line).

The optimal threshold is defined as the threshold maximizing the fraction of total variability explained (maximum of the blue line). As can be seen in Figure S2, our choice of threshold of 0.5 is relatively close to the optimum value that is 0.25 for the data set from [35], and 0.2 for the data set from [30]. Moreover, as Figure S2 illustrates, the different plotted statistics do not change markedly due to this sub-optimal choice of threshold and thus this choice does not affect our results significantly.

As different proteins have very different average fractions, the aforementioned calculation may be biased towards proteins with higher average fractions. To avoid this effect, the analysis presented was performed on the normalized fractions as defined in 4.1.1.

The noise in current whole proteome measurement techniques makes it difficult to distinguish between proteins that scale coordinately, as is predicted by our model, and proteins that scale differentially, but within measurement uncertainty. Thus, it is unclear to what extent the effect we predict affects actual protein fractions versus their possible individual up regulation with growth rate. We expect future improvements in the accuracy of whole proteome measurements to quantitatively reveal the importance of passive coordinated scaling with growth rate in shaping the proteome composition. These coming improvements in accuracy will enable better testing of the scope and validity of the model presented here.

**Figure S2:**
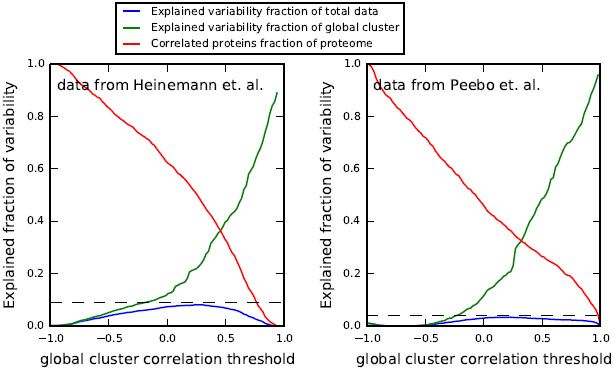
Statistics on the explained variability in the normalized data set as a function of the threshold used for defining strong correlation with growth rate. An optimal threshold is a threshold that maximizes the fraction of explained variability in the proteome by linear regression on proteins that have a correlation with growth rate that exceeds the threshold (blue line). The maximal explained variability in each data set is marked as a horizontal dashed line and is 0.082% (obtained given a threshold of 0.25) for the data set from [35], and 0.035% (obtained given a threshold of 0.2) in the data from [30].

### 6.3 Analysis of the data set from [35] including fast growth conditions

Due to the fast growth rate under LB and AA supplemented glycerol, compared with the other conditions measured in the data set from [35] these conditions were not included in our primary analysis as was noted in section 4.1.3. Including these conditions result in a much smaller set of proteins with a strong positive correlation with growth, as many of the proteins in that group in the slower conditions get down-regulated when AA are added to the media, significantly reducing their Pearson correlation with growth rate. For example, the Pearson correlation with growth rate of gapA, involved in glycolisys, drops from 0.73 to 0.35 when these conditions are included. Another such example is glyA, involved in serine and threonine metabolism, that has a correlation with growth rate of -0.12 when the faster conditions are included in the analysis vs. a correlation of 0.7 when they are excluded.

Figure S3 shows the implications of including the fast growth conditions in the analysis. As can be seen, many proteins are now less correlated with growth rate due to down regulation under the fast conditions. However, despite having fewer proteins being strongly positively correlated with growth (352 vs. 473) and despite the accumulated fraction of these proteins being lower under the slower growth conditions (≈ 30% vs. ≈ 40%), these proteins do occupy > 50% out of the proteome under fast growth in LB.

**Figure S3:**
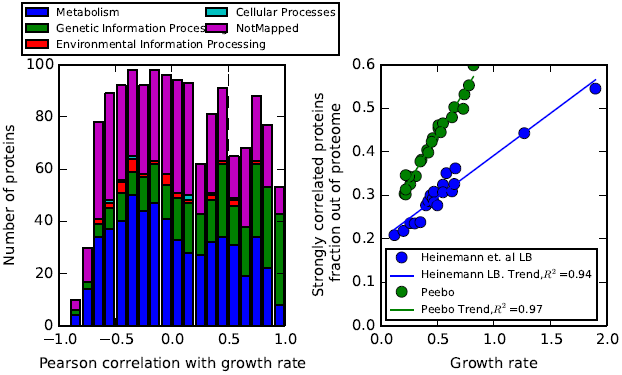
Including growth in LB and AA supplemented glycerol media in the analysis of the data set from [35]. Fewer proteins are strongly positively correlated with growth but these proteins form more than 50% of the proteome in fast growth.

### 6.4 Discussion of reasons for differing ribosome concentration relation to growth rate

Differences in ribosome concentration across growth rates as reported in different studies can result from a few factors:

1. Different growth rates and conditions monitored.
2. Inaccuracies and differences in the proteomic analysis procedures.
3. Usage of different strains.
4. In many studies the amount of ribosomes is deduced by measuring the RNA to protein ratio, assuming a relatively fixed portion of the RNA is rRNA. In our study, in contrast, ribosomal proteins are used as a proxy for estimating ribosomes concentration and, moreover, the RNA to Protein ratio is assumed to be constant. Therefore, and as it is known that ribosomes can operate even in the absence of some ribosomal proteins, such differences in manner of inference can account for some of the differences encountered.

### 6.5 The fraction out of the proteome of proteins that are not differentially regulated between conditions can be predicted by referencing other such proteins

**Figure S4:**
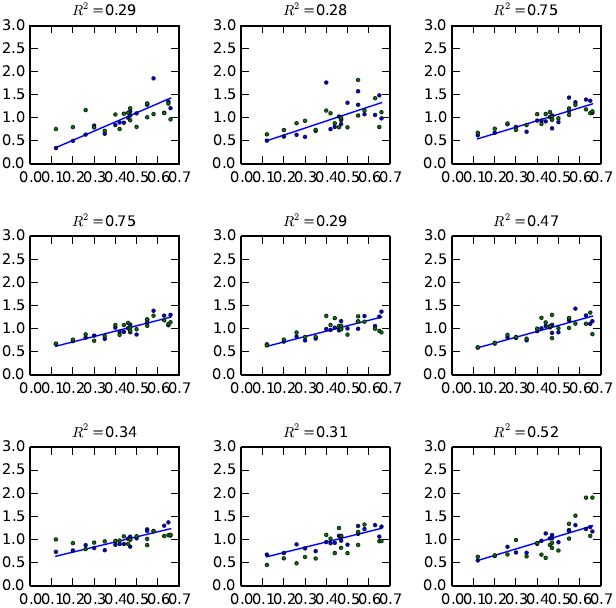
A selection of random predictions of protein fractions from the highly correlated with growth rate group, taken from the data set of [35]. Each panel shows the average fraction of 10 random proteins that are highly correlated with growth (blue dots), a regression line that best fits the data, and the fraction of a different random protein (green dots). The *R*^2^ value for the trend line and the different protein is given.

### 6.6 Breakdown by function of proteins strongly correlated with growth rate

**Table S1:**
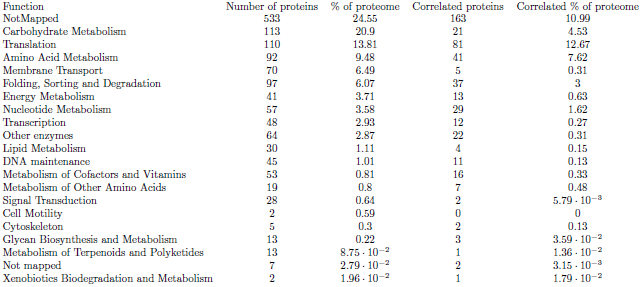
Breakdown by function of strongly positively correlated with growth rate proteins in the data set from [35]

**Table S2:** Breakdown by function of strongly positively correlated with growth rate proteins in the data set from [30]

| Function | Number of proteins | % of proteome | Correlated proteins | Correlated % of proteome |
| --- | --- | --- | --- | --- |
| NotMapped | 387 | 23.86 | 67 | 8.81 |
| Carbohydrate Metabolism | 104 | 20.58 | 19 | 4.11 |
| Translation | 97 | 17.65 | 79 | 17.05 |
| Membrane Transport | 65 | 8.89 | 5 | 0.24 |
| Amino Acid Metabolism | 81 | 6.27 | 20 | 1.23 |
| Folding, Sorting and Degradation | 82 | 5.01 | 23 | 1.86 |
| Energy Metabolism | 41 | 4.28 | 19 | 2.79 |
| Nucleotide Metabolism | 47 | 3.62 | 30 | 2.9 |
| Transcription | 33 | 2.58 | 7 | 1.36 |
| Other enzymes | 46 | 2.11 | 5 | $8.53 \cdot 10^{-2}$ |
| Lipid Metabolism | 18 | 1.29 | 5 | 0.59 |
| DNA maintenance | 33 | 1.14 | 5 | 0.17 |
| Metabolism of Cofactors and Vitamins | 39 | 0.72 | 8 | 0.25 |
| Metabolism of Other Amino Acids | 17 | 0.65 | 4 | 0.44 |
| Cell Motility | 5 | 0.43 | 1 | $4.98 \cdot 10^{-2}$ |
| Signal Transduction | 23 | 0.31 | 5 | $3.92 \cdot 10^{-2}$ |
| Cytoskeleton | 5 | 0.27 | 0 | 0 |
| Glycan Biosynthesis and Metabolism | 10 | 0.23 | 2 | $7.49 \cdot 10^{-2}$ |
| Metabolism of Terpenoids and Polyketides | 7 | $5.21 \cdot 10^{-2}$ | 1 | $1.51 \cdot 10^{-2}$ |
| Xenobiotics Biodegradation and Metabolism | 2 | $5.14 \cdot 10^{-2}$ | 0 | 0 |

### 6.7 Ribosomal proteins scale similarly to non-ribosomal proteins that are strongly positively correlated with growth rate

Comparing the normalized sum of ribosomal proteins to the normalized sum of the positively correlated with growth rate proteins that are non-ribosomal shows that these two groups scale in the same way with the growth rate, as is seen in Figure S5

**Figure S5:**
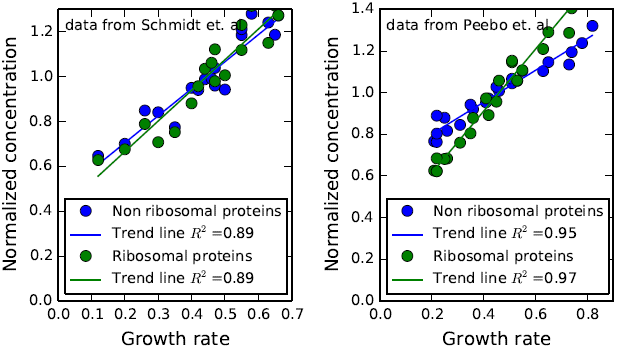
The scaling with growth rate of ribosomal proteins and non-ribosomal, but highly correlated with growth rate proteins is shown. Trend lines for the two groups of proteins are plotted. The scaling with growth rate is similar between the two groups of proteins.

### 6.8 Additional figures of simulated and randomized data sets

The maximal explained variability in data sets with shuffled protein abundances is significantly smaller than in the real data sets as is seen in figure S6.

A simulated data set, assuming half of the proteins scale linearly with growth rate with normalized intercept at 0.5, similar to the intercept found in the data analysis, and with simulated normally distributed noise levels of 25%, result in distributions similar to those found in the original data analysis (Figure S7)

**Figure S6:**
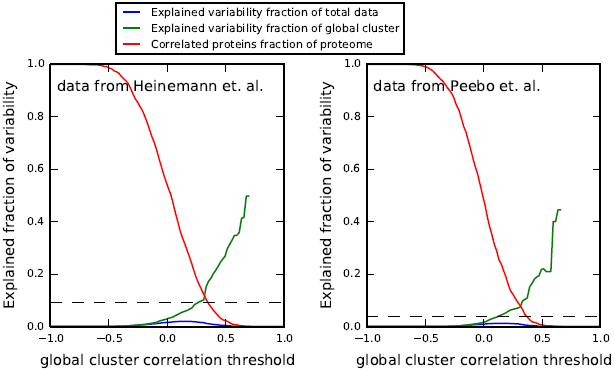
Fraction of explained variability by linear regression on the group of strongly positively correlated with growth rate proteins for the shuffled data sets.

**Figure S7:**
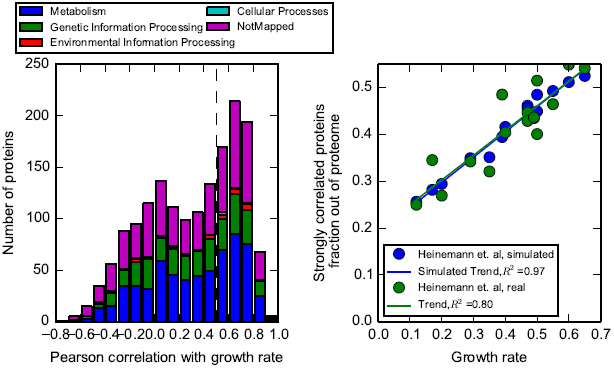
A simulated data set, assuming half of the proteins are perfectly correlated with growth rate and half are fixed, with simulated noise level of 25%. Average protein fractions, growth rates and normalized slope of the correlated proteins are based on the data set from [35]. The normalized intercept of the correlated proteins was set to 0.5 in accordance with the intercept found in the original data analysis. The results are similar to those obtained for the real data set, showing that, given the experimental noise, identical coordination with growth rate of half of the proteins would result in similar outcomes to those observed in the data sets we use.

